# Changeover from signalling to energy-provisioning lipids during transition from colostrum to mature milk in the giant panda (*Ailuropoda melanoleuca*)

**DOI:** 10.1101/063701

**Authors:** Tong Zhang, David G. Watson, Rong Zhang, Rong Hou, I. Kati Loeffler, Malcolm W. Kennedy

**Affiliations:** Strathclyde Institute of Pharmacy and Biomedical Sciences, 161, Cathedral Street, Glasgow G4 0RE, Scotland, UK; Institute of Clinical Pharmacology, Guangzhou University of Chinese Medicine, No. 12 Jichang Road, Guangzhou 510405, P.R. China; Sichuan Key Laboratory of Conservation Biology for Endangered Wildlife, Chengdu Research Base of Giant Panda Breeding, 1375 Panda Road, Northern Suburb, Chengdu, Sichuan Province 610081, P.R. China; Institute of Biodiversity, Animal Health and Comparative Medicine, and Institute of Molecular Cell and Systems Biology, College of Medical, Veterinary, and Life Sciences, Graham Kerr Building, University of Glasgow, Glasgow G12 8QQ, Scotland, UK.

## Abstract

Among the large eutherian (‘placental’) mammals, ursids (bears) give birth to the most altricial neonates with the lowest neonatal:maternal body mass ratios. This is particularly exemplified by giant pandas in whom the transition from colostrum to main-phase lactation is unusually prolonged. To examine whether there is compensation for the provision of developmentally important nutrients that other species groups may provide in utero, we examined colostrum and milk lipids from birth until the transition was complete. Lipids known to be developmental signals or their precursors, and those that are fundamental to nervous system construction, such as docosahexaenoic acid (DHA) and phosphatidylserines containing DHA, appear early and then fall dramatically in concentration to a baseline at about 20 days. This also applies to other signalling lipids such as lysophosphatidylserines. The dynamics of lysophosphatidic acid and eicosanoids display a similar pattern, albeit less clearly and with differences between mothers. Triglycerides occur at relatively low levels initially and then increase in concentration with time post-partum until a plateau is reached at about 30 days or later. These patterns indicate an early provision of signalling lipids and their precursors, and lipids crucial to brain, retinal and central nervous system construction, followed by a changeover to lipids for energy metabolism. Thus, in giant pandas, and possibly among ursids in general, lactation is adapted to provisioning a highly altricial neonate to a degree that approximates to an extension of gestation.

**Abbreviations**
AA: arachidonic acid
ALA: alpha-linolenic acid
DHA: docosahexaenoic acid
EPA: eicosapentaenoic acid
FFA: free fatty acid
GL: glycerolipid
HPLC: high-performance liquid chromatography
HRMS: high resolution mass spectrometry
LA: linoleic acid
LC: liquid chromatography
LL: giant panda Li Li
lysoPS: lysophosphatidylserine
MS^1^, MS^2^or MS/MS MS^3^, MS^n^: multiple stage ion fragmentation mass spectrometry
OPLS-DA: orthogonal partial least square-discriminant analysis
PC: phosphatidylcholine
PE: phosphatidylethanolamine
PI: phosphatidylinositol
PS: phosphatidylserine
PUFA: polyunsaturated fatty acid
SM: sphingomyelin
TAG: triacylglyceride
XYT: giant panda Xiao Yatou
YY: giant panda Yuan Yuan

## Introduction

Milk is the sole source of nutrition for mammalian neonates, and is also an essential conduit of immune support for many species of infant. Mammary secretions change dramatically from colostrum (‘first milk’) to mature, main-phase milk during the immediate post-partum period. Broadly, colostrum tends to be more protein-rich and lipid-poor than later milk, and is particularly rich in immunoglobulins and innate anti-microbial factors [1–7]. The change to mature milk represents a switch to more energy-rich nutrition in which sugars and fats may predominate [1, 2].

The neonates of some species are absolutely dependent for their survival and development on colostrum from their mothers, whilst others are less so. The period of time for this dependence – and hence the duration of the colostrum phase of lactation – also varies among species. The difference is largely a function of the type of placenta involved, where, for instance, species with epitheliochorial placentae cannot transfer immunoglobulins from maternal to foetal circulations (as in ungulates), and in whom colostrum is thereby essential to survival of neonates [2]. At the other extreme are species with haemochorial placentae (e.g. humans) whose placentae transport immunoglobulins from maternal to foetal blood circulations prior to birth, although this is confined to immunoglobulin G (IgG) [2–4].

We are investigating the lactation strategies of ursids (bears) because, amongst eutherian (‘placental’) mammals, they give birth to the most altricial (developmentally immature) neonates with the lowest neonate:maternal body mass ratios [8]. This may mean that lactation is adapted in two main ways to provide for such immature neonates. First, their milks may change more slowly from colostrum to main-phase than in other species. Second, the components of early ursine colostrum may be unusual.

We chose giant pandas for this study because they exhibit an extreme amongst ursids in the altriciality of their neonates [9]. In addition, the captive breeding program for giant pandas in China and its associated intensive human handling of the animals provided an opportunity for serial milk sampling in the immediate post-partum period. In our previous studies of panda milk, we discovered the transition between colostrum to mature-phase milk to be unusually, perhaps uniquely, prolonged amongst eutherians [10]. That study indeed revealed a slow maturation in the protein profiles, the changeover and maturation process taking approximately thirty days to complete, over which period certain species of oligosaccharide disappeared, whilst others appeared. Further detail was added in a second study, in which a broad-spectrum metabolomics approach discerned three phases in the transition from colostrum to mature lactation [11]. The milks of different mothers in that study were at first similar in composition but then diverged after about seven days.

There are no precise criteria that define the end of the colostral period and the onset of main-phase lactation for any species. We here take the colostrum phase to end when all the major components of milk reach an approximate steady state, although slight modifications in composition may still occur during the main phase. By this definition, the colostrum period of giant pandas ends at about 30 days [10, 11], which is considerably longer than is known for any other Eutherian, and stands in stark contrast to the less than 24 hours in, for example, grey seals (A. Lowe, P. Pomeroy, D.G. Watson, M.W. Kennedy, unpublished).

Lipids in milk could be broadly divided into those needed for metabolic energy support, the construction of membranes and antimicrobial activities, and those that serve as immune and developmental signalling molecules or their precursors. Energy for foetuses is almost exclusively provided in the form of trans-placental delivery of glucose and lactate [12]. Triglyceride transfer for this purpose is minimal or zero, and foetal gluconeogenesis is essentially inactive [12]. After birth, however, milk offers a switch to lipid-and lactose-based energy provision, the balance of which varies from species to species [1, 2].

Large-brained mammals exhibit a heavy requirement for polyunsaturated fatty acids (PUFAs) for brain development [12, 13]. This appears to be satisfied in utero by translocases that transport crucial PUFAs such as arachidonic (AA) and docosahexaenoic (DHA) acids en masse from the maternal to the foetal circulation, resulting in higher levels of PUFAs in foetal than maternal circulations [12]. Trans-placental transportation of PUFAs may be confined to species with haemochorial placentae [12], is minimal in those with epitheliochorial placentae (e.g. sheep; [14]), and may also be for species with endotheliochorial placentae [15]. The latter placental type is widespread within the Carnivora [16–18] so there may therefore be a need for immediate post-partum provision of PUFAs for continued neonatal development that may be particularly urgent for highly altricial neonates.

Here we show that there is a slow transition between colostrum and mature milk lipid profiles in giant pandas, and that some species of lipid are at first abundant and then fall away, whilst others increase with time and stabilise in concentration. The main changeover, like that which we demonstrated for the protein, oligosaccharide and metabolome profiles, occurs between twenty and thirty days. Importantly, those lipids such as PUFAs that are considered to be crucial for developmental and immune signalling and for brain growth are present at their highest levels only very early in lactation, while those needed predominately for energy generation and membrane construction appear later.

## Results

### Lipid profile of giant panda milk

The analysis of milk lipids required sequential rounds of mass spectrometry and ion fragmentation, which was particularly required for monoacyl and diacyl glycerolipids in which the acyl tails are heterogeneous. Added to which is the complexity of the positions of carbon:carbon double bonds. A summary of the main types of lipids that were identified is given in Table 1. (A more detailed version of this information is provided in Supplementary Table S1, which contains more indicators of how the different classes yielded ion fragments.) The principal lipids were phospholipids, sphingomyelins, GLs (triacylglycerols), free fatty acids (FFAs) and sterols, with a few other classes contributing to a total of 403 distinct species. The lipids are typical of mammalian milks, and ranged as expected from those broadly involved in energy provision (TAGs), membrane construction (phospholipids), precursors of developmental and immune or inflammatory signals, and lipids enriched in brains (PUFAs). Statistical analysis showed that these different lipid classes had distinct patterns of increase and decrease in the milk with time after parturition (Figures 3 and Supplementary Figure S2).

**Table 1.** Summary of lipids found in giant panda milk.

| Main class | Subclass | Number of species detected | Examples of deductions from multiple stage mass spectrometry (MS<sup>n</sup>) analysis. (C:D)<sup>a</sup> |
| --- | --- | --- | --- |
| Phosphatidylserine (PS) | Monoacyl-PS | 25 | PS (18:0/0:0) |
|  | Diacyl-PS | 38 | PS (18:0/22:6) |
| Phosphatidylcholine (PC) | Monoacyl-PC | 23 | PC (18:1/0:0) |
|  | Diacyl-PC | 43 | PC (16:0/16:0) |
| Phosphatidylinositol (PI) | Monoacyl-PI | 2 | PI (20:4/0:0) |
|  | Diacyl-PI | 16 | PI (20:4/18:0) |
| Phosphatidylethanolamine (PE) | Monoacyl-PE | 16 | PE (18:1/0:0) |
|  | Diacyl-PE | 19 | PE (18:1/16:0) |
| Sphingomyelin (SM) |  | 27 | SM (d18:1/24:1) |
| Glycerolipids (GL) |  | 49 | Triacylglycerol (TAG) (52:6) |
| Free fatty acid (FFA) |  | 23 | Docosahexaenoic acid (DHA) (22:6) |
| FFA derivatives |  | 65 |  |
| Sterols |  | 14 |  |
| Others |  | 43 |  |
| **Total** |  | **403** |  |

<sup>a</sup> C, number of carbon atoms; D, number of double bonds.

### Sequential changes and progressive divergence between individual mothers

Figure 1 shows the score plot of the OPLS-DA model generated by comparing the samples collected before (red spots) with after (blue spots) 7 days postpartum. Samples collected from all three mothers before day 7 show similar rates of change with time. Their clustering around zero on the vertical axis indicates that their compositions are similar to one another. In contrast, the wider scatter on the vertical axis after day 7 indicates progressive, rapid and approximately simultaneous divergence between individual mothers, as noted previously for other milk components [11].

**Figure 1.**
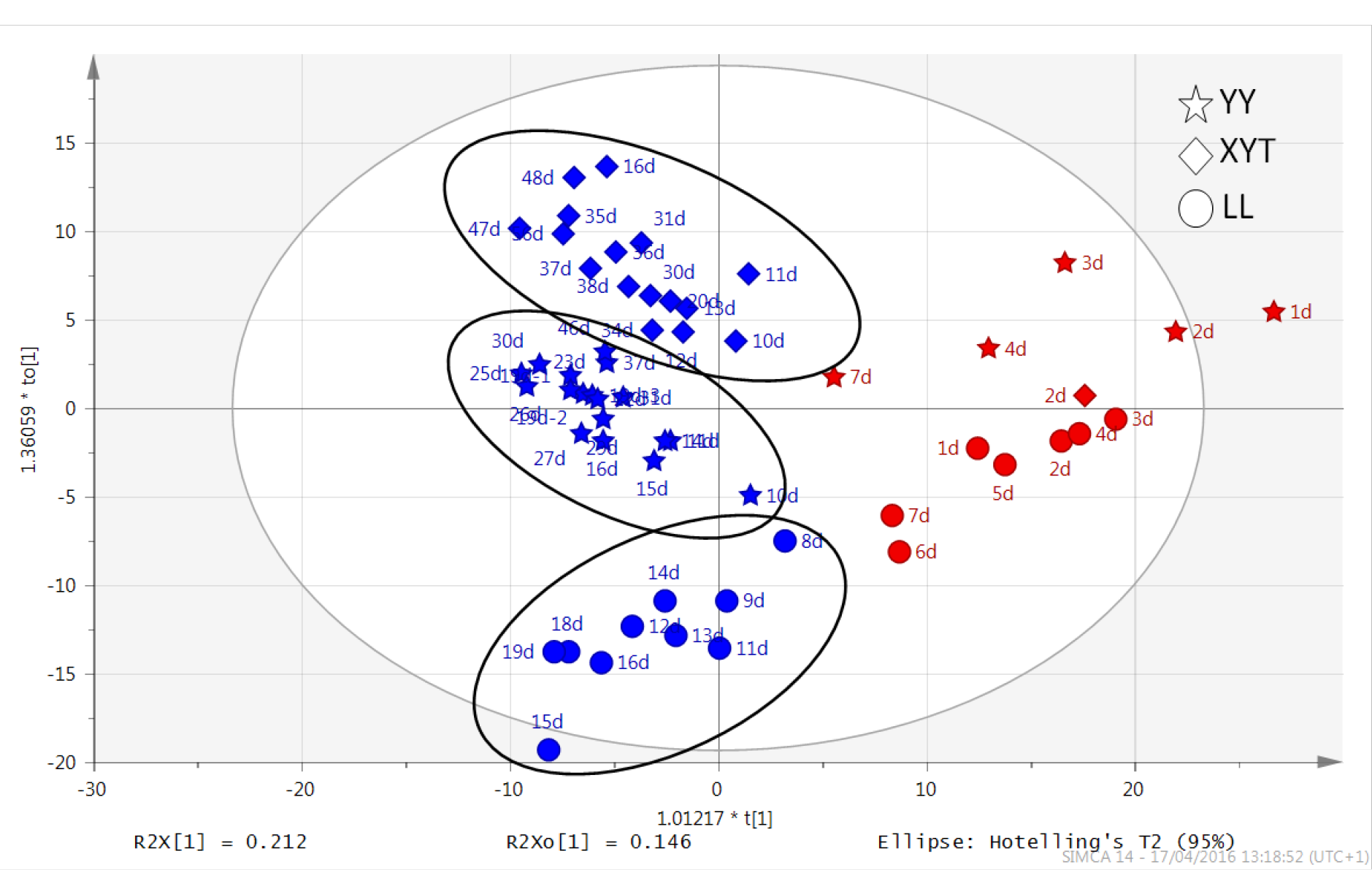
OPLS-DA score plots of milk samples collected serially from three giant pandas after birth. Data from 55 samples collected from Yuan Yuan (YY; stars), Xiao Yatou (XYT; diamonds), Li Li (LL; circles). This shows that the lipid profiles of early colostrum (red) are similar between individual mothers, though with indications of progressive divergence with time after birth, followed by accelerating divergence from about 7 days (blue). The changes in concentrations of selected classes of lipids are illustrated in Figure 3, and in Supplementary Figures S1 and S2.

Figure 2 shows the distribution of the lipid profiles as an S-plot from the OPLS-DA model. Data points located far out on diagonally opposite wings of the “S” indicate lipids that changed significantly and consistently in relative abundance from colostrum to mature milk. The four lipids at the most extreme positions on the “S” distribution (labelled in Figure 2) were identified as choline phospholipids (PCs) but with different acyl chain compositions.

**Figure 2.**
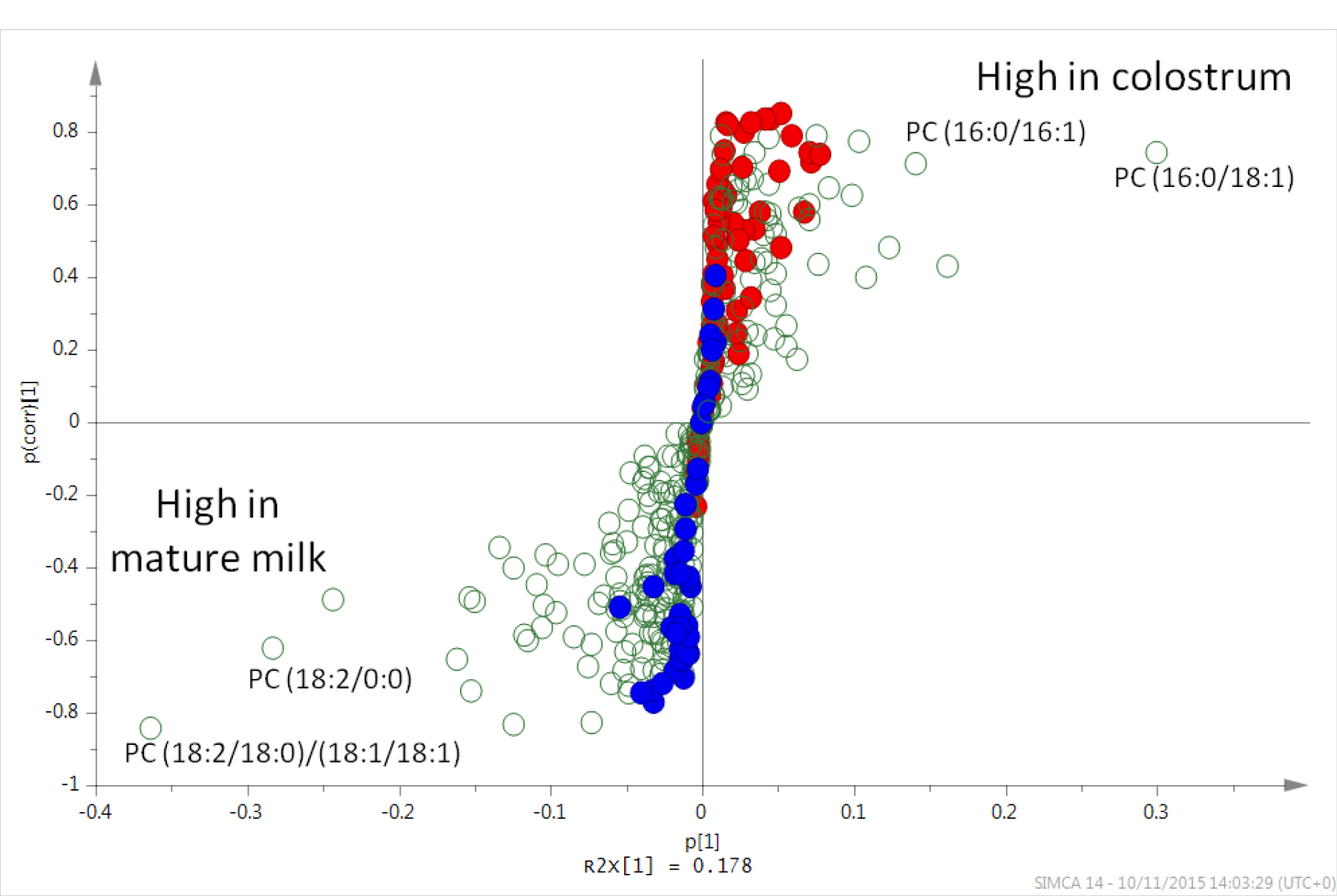
Changes in giant panda milk lipids in early lactation. Each point represents a single lipid species. Those appearing in the top right arm of the S were high in colostrum and low in mature milk, and those to the bottom left arm were most abundant in mature milk and relatively low in colostrum. This S-plot derives from orthogonal partial least squares discriminant analysis (OPLS-DA) of 55 giant panda milk samples collected serially after birth from three giant pandas. The x-variable is the relative magnitude of a variable which reflects the abundance of the lipid in panda milk to some degree (see the Supplementry for a more detailed explanation). The y-variable is the confidence/reliability such that a positive value for a lipid indicates that it is high in colostrum (to a maximum of +1); a lower and to a negative value indicates that that lipid is low in colostrum and increasingly high in mature milk (to a minimum of −1). Values in the centre of the plot near 0 are close to the noise level, representing a high risk for spurious correlations, and therefore appear to be constant between colostrum and main phase milk. Thus, data points falling in the upper right or lower left corners of the plot represent those whose representation of discrete milk phases may be deduced with the greatest confidence. GLs are shown in blue, PS lipids in red, and other lipid classes (including PCs) in open circles. The changes in relative concentrations with time after birth for each of the four compounds named in the diagram are plotted in Figure 3.

**Figure 3.**
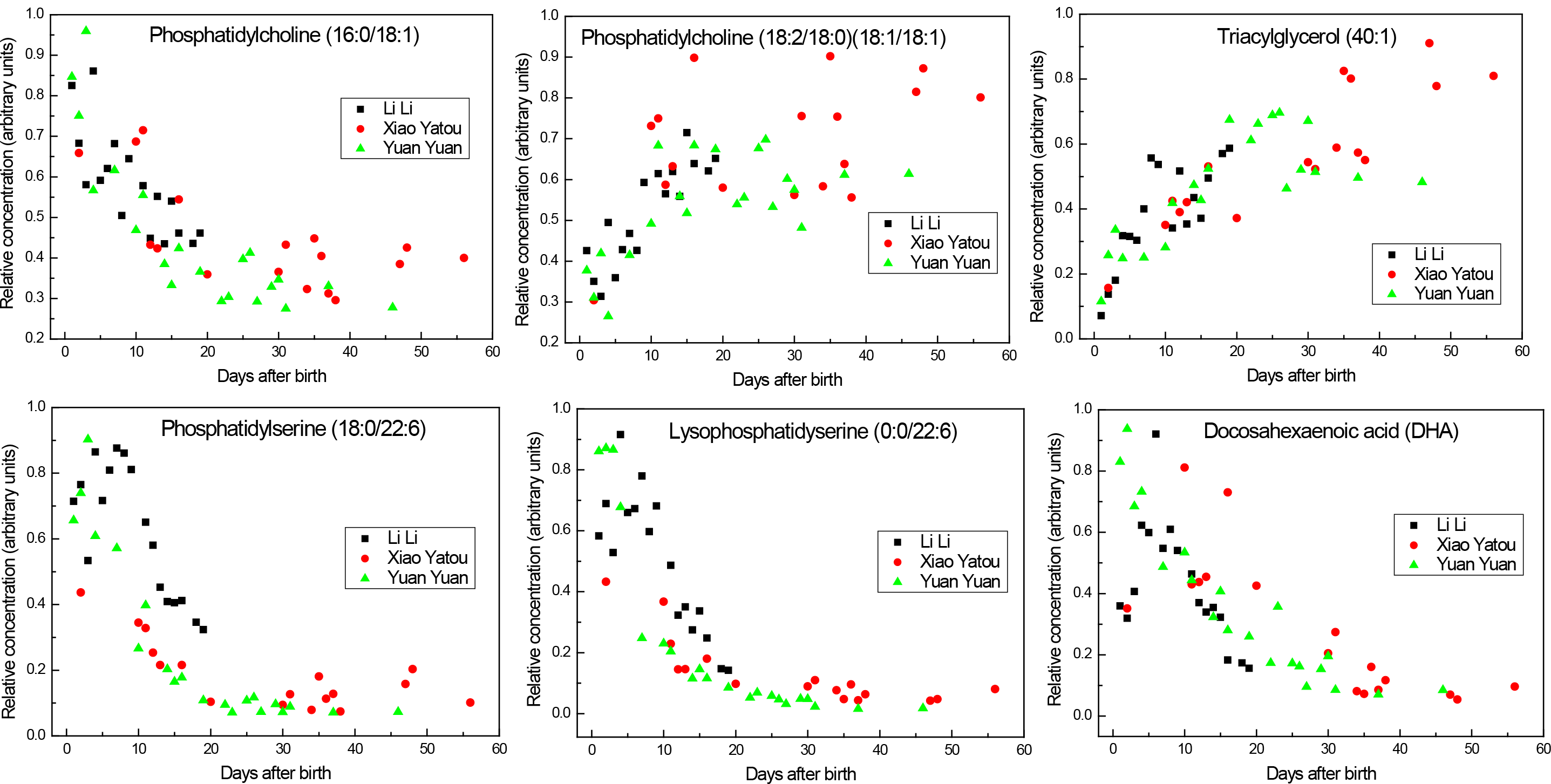
Time course of post-parturition changes in selected lipids in giant panda milk. The values are relative areas under liquid chromatography signal peaks for each lipid. These relative values show changes with time after birth for each lipid species, and are not comparable between the different lipid species.

These lipids exhibited divergent concentration trends during the lactation period such that PC (16:0/18:1) was initially abundant in colostrum whereas the (18:2/18:0) (18:1/18:1) isoform only became abundant in mature milk. Plots of the relative concentrations of these two lipids with time show that the changeover was complete by about day 20 (Figure 3).

Other lipid classes, however, showed either no significant change or a weaker differentiation between colostrum and mature milk than did the PCs. For instance, in Figure 2 the GLs (indicated in blue) distribute along the vertical axis from 0.4 to −0.8 (low in colostrum but high in mature milk), and phosphatidyserines (PS; in red) had a slightly overlapping distribution along the y-axis from −0.3 to 0.85 (high in colostrum but low in mature milk). In both cases, the data points cluster close to the 0 on the horizontal axis, indicative of either low abundance, or, more likely, weaker MS signals than other compounds because they ionise poorly. Interestingly, the time course for PSs was also followed by lysoPSs in being high in colostrum and low in mature milk (Figure 3).

The level of TAGs dramatically increased with time postpartum, reaching a plateau after about 30 days (Figure 3). The kinetics of this change was similar for all three giant panda mothers. The increase in TAGs mirrors a cruder observation of a change in fat content with time in that the low density fat layer appearing on milk samples that were centrifuged in the cold is small in colostrum and increases with time (Supplementary Figure S1).

### Developmental and signalling lipids and their precursors

Of particular interest, given their roles as precursors to pharmacologically and immunologically active compounds, certain PUFAs (e.g. DHA, Figure 3) are relatively abundant very soon after birth but then decay to a low plateau by about 30-40 days. The same applied, albeit less clearly, to eicosapentaenoic (EPA) and arachidonic acids (AA) (Supplementary Figure S2). Linoleic (LA) and α-linolenic acids (ALA), both of which are absolute essential fatty acids for humans, increased with time after birth (Supplementary Figure S2).

A group of lipids that is increasingly recognised as being important in signalling is the PSs. Additional multiple stage MS analysis of the isoforms in the milk samples showed that the acyl chains of the major PS components comprised 16, 18, 20 or 22 carbons with a diverse degree of unsaturation from 1 to 6 C=C bonds. Unlike other polar lipids (e.g. PCs) some PSs were detected with highly unsaturated acyl chains, e.g. PS (18:0/22:6) and lysoPS (0:0/22:6) (Figure 3, Supplementary Figure S3), which we have not found to have been reported in studies of polar lipids in other milks using similar analytical techniques.

Figure 3 also shows the development of these two PSs during early lactation, showing that their levels are high in the first 7 days then decline rapidly (for one of the animals, Yuan Yuan, in a close to exponential decay) to a low plateau by about 20 days postpartum. Interestingly, a similar time course was obtained with DHA, which is also present as the unsaturated acyl chain of these two PSs (Figure 3). The neonate is therefore supplied with this conditionally essential fatty acid either as the free fatty acid (FFA) or as one chain of the predominant PSs.

Lysophosphatidic acid, which also has several direct biological activities, showed a distinct exponential decline with time after birth in one animal (again, Yuan Yuan; Supplementary Figure S2), but there was only slight evidence of this for Li Li (for whom the sampling period was short), and none for Xiao Yatou.

## Discussion

The founding question to this work was whether the bearing of highly altricial neonates by ursids is mirrored by a lactation strategy that is unusual amongst eutherians. Specifically, is there a prolonged transition period between colostrum and main-phase milk production, and are the constituents of the milk modified to compensate for the early developmental state of the neonates?

In the ursid species that exhibits the highest degree of altriciality, the giant panda, we have found that there is indeed a prolonged maturation phase in lipid profiles that takes about thirty days to complete. This parallels the time course for other components of giant panda milk, namely proteins and oligosaccharides [10], and the overall metabolome [11]. The changes in lipid profiles reveal a dramatic shift in the functional balance of the lipid content from an early predominance of lipids that are precursors of signalling molecules and that are essential for construction of the central nervous system, to those predominately involved in energy metabolism.

### Signalling lipids dominate the lipid profile of the prolonged colostrum period

Mammals are unable to introduce C=C double bonds in fatty acids beyond carbons 9 and 10, which is why the polyunsaturated linoleic acid (LA; ω-6; 18:2, 9, 12; 18:2n-6) and α-linolenic acids (ALA; ω-3; 18:3, 9, 12, 15; 18:3n-3) are essential in the diet. The so-called conditionally essential fatty acids are derivatives of these and include DHA, AA, and EPA. While some of these PUFAs are major components of neuronal tissues, they also have central roles in developmental and immune activation as precursors to signalling lipids such as prostaglandins and leukotrienes. The lipid classes that predominate early and then diminish in giant panda lactation include DHA, both as the free fatty acid and as an acyl chain of a PS and a lysoPS, all of which show similar time courses (Figure 3). DHA is crucial to visual acuity and neural development and is the most abundant ω-3 fatty acid in the brain and retina of mammals [13, 19–25]. DHA also modulates the carrier-mediated transport of choline, glycine, and taurine [26], the latter being crucial to neuronal development [27], and is possibly a dietary-essential compound in bears as it is in some other Carnivora [28–30]. The time courses for the decline of two other PUFAs, AA and EPA, in colostrum are similar to that for DHA though less well defined (Supplementary Figure S2).

The decline in DHA in giant panda milk with time, which is also seen in humans colostrum [31], could be for several reasons. First, that lipid desaturation mechanisms in neonates required to convert LA and ALA may initially be inadequate in highly altricial neonates (although this does not apply to preterm humans [32]). Second, the substantial requirement of neonatal mammals for DHA may not be sufficiently supplied by the relatively low absolute fat content of colostrum and elevated levels of the free fatty acid are supplied to compensate [31]. Third, the intestines of neonatal mammals are susceptible to inflammatory conditions that may be ameliorated by anti-inflammatory lipid mediators such as resolvins derived from DHA (and EPA), potentially explaining lower incidence of intestinal inflammation in breast-fed rather than formula-fed human infants [31]. At this stage our analysis has not discriminated the anti-inflammatory resolvins and lipoxins from other eicosanoid lipids, although we did find diminishing levels of the pro-inflammatory leukotrienes (Supplementary Figure S2) such as are also found in human milk [31].

Other important roles of milk fatty acids include direct anti-microbial activities [33] that will be relevant to a newborn suddenly exposed to a microbe-rich environment and whose gut microbiome is naïve and must establish a functional balance. Of particular potential importance to a hairless, altricial neonate such as a giant panda cub, is the integrity of the epidermal permeability barrier to avoid transcutaneous water loss. There is a particular role for linoleic acid in the regulation of epidermal water-permeability, in the form of special O-acylated ceramides ([34–38]).

PSs comprise important membrane structural components in most cells. They are particularly enriched in the inner leaflet of the plasma membrane in neural tissues where they are key to several signalling pathways and make up 13-15% of the phospholipids in the human cerebral cortex [39, 40]. Moreover, PS-dependent signalling, like DHA, is directly involved in processes of neuronal differentiation and survival [39, 40]. These roles further emphasise the potential significance of maternal delivery of PS, lysoPS and DHA in the immediate neonatal period (Figure 3). The PS receptor, and therefore also its ligand, is essential during embryogenesis in mice, which also produce altricial neonates [41].

PSs also form protein-phospholipid complexes that initiate calcification during the formation of bone [42, 43], which again could be particularly relevant to a highly altricial neonate. The lyso forms of PSs are particularly notable for involvement in cellular signalling, including neural development, that could be crucial to development of mammalian embryos and survival of neonates [44–46]. Also potentially crucial to a newborn is that lysophospholipids, especially lysoPSs, are increasingly being found to have potent immunoregulatory properties [44, 47–52].

The other major class of phospholipids in giant panda milk, the PCs, are major components of biological membranes and pulmonary surfactant. Our analysis of giant panda colostrum indicated changes in provision of PCs to the neonate with time (Figure 3). There was, for instance, an intriguing complementarity in time course over the first 20 days, in which one PC isoform (16:0/18:1) was present early and then diminished, whereas the abundance of another (18:2/18:0)(18:1/18:1) was initially low and then increased (Figure 3).

### Triacylglycerides appear in mature milk

Lipids that dominate the profile of early colostrum appear to be those involved in a range of developmental functions. In contrast, TAGs, which are likely to be more involved in provision of energy, appear only slowly with time (Figure 3). An early deficit in energy supply may be associated with the slight loss of body mass seen in panda cubs (as is common amongst mammal neonates) immediately after birth [11], but also that neonatal cubs may not be able to digest TAGs efficiently. The persistence of bile-salt-activated lipase (commonly found in milks and thought to aid digestion of triglycerides) in giant panda milk supports this suggestion, in that their maternal provision may be necessary until a cub’s own pancreatic production of the enzyme becomes sufficient [10].

With regard to energy provision, it is also noteworthy that lactose is present in giant panda milk initially but rapidly diminishes with a time course that is the inverse of that for TAGs ([10, 11] and Figure 3). A similar, though much slower conversion from sugar-to lipid-based energy supply occurs in marsupials [53, 54].

The imaginative hypothesis [8] explaining why ursids produce such altricial neonates states that mobilisation of fat reserves for gluconeogensis to supply embryos trans-placentally with glucose is metabolically wasteful during hibernation. Birthing at an early stage of development allows a switch to lipid-based energy supply via milk that reduces the metabolic demand on a hibernating mother. However, giant pandas, like several other species of ursid (Andean, sloth and sun bears), do not hibernate and there is no indication yet that they had ancestors that did so.

It is nevertheless clear from this and our previous studies [10, 11] that the altriciality of giant panda cubs is indeed reflected in a prolonged maturation from colostrum to main-phase lactation which takes about twenty days for lipids that are important in signalling and developmental functions (this study) and about thirty days for TAGs (this study), protein and oligosaccharide components [10, 11]. Within these transitions, there are intriguing sublteties in the changeovers between compounds, such as for the two PC isoforms mentioned above, and oligosaccharides in which early-and late-appearing forms differ merely in their glycosidic linkages [10].

### Artificial milk replacers cannot mimic the composition and dynamics of neonatal ursine requirements

Whilst our main interest is in lactational biology of ursids as a whole, the dramatic, time-dependent changes in the lipid content of giant panda milk and the associated complexity of biochemical composition emphasise the inadequacy of artificial milk formulae that are used to supplement or replace giant panda milk. As shown in Supplementary Figure S3, none of three milk replacers used for giant panda cubs (including one recently developed specifically for giant pandas; [55]) provide any of the lipids that predominate for the first twenty days, this being particularly so for PUFAs such as DHA. The analysis presented here and in our proteomic and metabolomic analysis of giant panda milks [10, 11] indicates how critical the biochemical constitution and dynamic of milks are for optimal neonatal development of organ systems (the central nervous system in particular), the immune system, and for the cub’s innate immunity.

This draws into question the practice of feeding artificial milk formulae as a matter of routine, and the extent to which such a practice compromises particularly the neurological and immune development of giant panda cubs. Moreover, our previous studies have highlighted the risk that these artificial formulae pose to the health of cubs, such as with their high relative content of lactose [10, 11]. Taken together, these findings disagree with the simplistic approach for development of panda milk replacers [55] and emphasise instead the need to revise husbandry conditions to eliminate artificial feeding in breeding centres.

### Is ursine colostrum a way of supporting a form of external gestation?

The term “external gestation” was originally applied to marsupials because they produce neonates of extraordinary altriciality following a very short gestation, and exhibit dramatic alterations in milk components with time [54, 56, 57] that may serve to provide nutrients required for early development that eutherians provide in utero.

Specifically with regard to lipids, one of the most dramatic changes in marsupial milks is exhibited by a family of proteins, the lipocalins, that is widely associated with transport of small lipids [58–60]. Changes in these proteins in marsupial milks may represent a changing requirement for lipid types, although there is currently no information on whether these different lipocalins exhibit discrete lipid transportation repertoires. While these marsupial milk-specific lipid carrier proteins do not occur in eutherians (though two forms of a lipid-transporting lipocalins, beta-lactoglobulin, do occur in giant panda milk; [10]), the changes in lipids we observed in the milk of giant pandas may indicate a similar role of milk in the provision of developmentally-essential lipids to an altricial neonate that in other species of eutherian are provided pre-partum. In this case, the term “external gestation” may indeed be applicable to bears.

## Materials and Methods

### Milk collection and processing

All giant panda samples were collected from captive-bred animals at the Chengdu Research Base of Giant Panda Breeding, Chengdu, Sichuan Province, P.R. China, during the years 2006, 2007, 2011, and 2012. Please see ref. [10] for a complete list of sampling dates, studbook numbers, and reproductive history of the individual animals. Milk was obtained by hand from trained, food-rewarded, conscious animals who were not anaesthetised, sedated, drug-treated, or physically restrained. The health status of each animal was monitored regularly, and while overt disease was not observed in any donor, minor ailments and suboptimal blood factor levels were observed in some. Mother pandas normally enter a period of anorexia for 7-10 days postpartum, though the animals from whom samples were obtained were given glucose before normal diet resumed. Milk samples were stored either in liquid nitrogen or at −80°C immediately after collection, transferred to Glasgow frozen, and stored at or below −20°C until use.

### Chemicals and standards

High-performance Liquid Chromatography (HPLC) grade acetonitrile and isopropanol were obtained from Fisher Scientific, UK. Ammonium formate was purchased from Sigma-Aldrich, UK. HPLC-grade water was produced by a Direct-Q 3 Ultrapure Water System from Millipore, UK.

### LC-HRMS/multiple tandem HRMS analysis and data processing

The preparation of milk samples and the high resolution mass spectrometry HRMS settings were described in our previous study [11]. In order to improve the separation of lipids, a silica gel column (ACE SIL, 150 × 3mm, 3μm, HiChrom, UK) was employed with the mobile phase A as a mixture of water and isopropanol (v/v 8:2) containing 20 mM ammonium formate, and phase B as acetonitrile and isopropanol (v/v 8:2). The LC gradient was programmed as follows: 90% of B from 0 to 5 min, decreasing to 70% at 9 min, 65% at 13 min, 60% at 23 min, 55% 28-30 min, and finally increased back to 90% at 31 min, which was then held until 40 min.

Peak extraction and alignment of the LC-HRMS data was carried out using MZMine 2.10 as previously [11]. As shown in Supplementary Table S1, different adduct forms were selected for annotation of different lipid classes and then searched on accurate mass against the database downloaded from the LIPID MAPS Lipidomics Gateway, http://www.lipidmaps.org/. Only putatively identified signals were used for subsequent statistical analysis.

Data-dependent multiple stage mass spectrometry (MS^n^) fragmentation scan was carried out with collision-induced dissociation at 35 V using a Surveyor HPLC system combined with a LTQ-Orbitrap mass spectrometer (Thermo Fisher Scientific UK) and the chromatographic conditions described above. The MS/MS spectra of some of the major lipids are gathered at the end of the Supplementary, and the interpretations are detailed in Supplementary Panel S1 and summarised in Supplementary Table S1.

### Statistical analysis

SIMCA version 13.0 (Umetrics, Umeå, Sweden) was used for multivariate analysis:Orthogonal Partial Least Squares Discriminant Analysis (OPLS-DA). The data were mean centred and Pareto scaled in order to generate an S-plot for visualisation of the components with significant influence in the dataset. The plots of peak areas of individual lipids versus days after birth were generated using Microcal ORIGIN software.

**Ethics statements** Milk sampling from giant panda mothers was carried out under ethical approval from the University of Glasgow's College of Medical, Veterinary & Life Sciences Ethics Committee, and the Chengdu Research Base of Giant Panda Breeding where the animals were held. All the giant panda mothers and their cubs were captive-bred and -maintained. Milk samples were collected from conscious, unrestrained animals trained to allow milk sampling during routine health checks and veterinary inspection, or when considered necessary to provide milk for orphan or abandoned cubs, or for research purposes. Handling of cubs during the milk collection period for health, normal growth, and veterinary checks is carried out daily at the Chengdu facility. Giant pandas are classified as endangered animals by the International Union for the Conservation of Nature (IUCN Red List 3.1). International transfer of milk samples from captive giant pandas in China to the laboratories in Scotland was covered by the CITES convention through permits issued by both donor and recipient countries. A permit was obtained from the Scottish Executive for the importation of the milk samples into Scotland as veterinary-checked animal products.

## Acknowledgements

We are grateful to the following at Chengdu Research Base of Giant Panda Breeding: Hairui Wang for supervising the collection and storage of the milk samples, Liang Zhang for administration of export arrangements and collection of veterinary records, and the Director, Professor Zhihe Zhang, for his continued support and encouragement of this project. The analytical work was funded by the University of Glasgow and the University of Strathclyde. The funding of mass spectrometry equipment at the University of Strathclyde was provided by the Scottish Life Sciences Alliance.

## Author contributions

Original concept and design of the study: MWK, IKL, RH.

Performed the analysis: TZ, RZ, DGW.

Analyzed the data: TZ, RZ, DGW, MWK.

Contributed reagents/materials/analysis tools: TZ, DGW, HR, HW, MWK.

Wrote the paper: MWK, TZ, DGW, IKL.

